# An Interactive Health Game Using Machine Learning: A Prototype

**DOI:** 10.1101/2020.12.01.405852

**Authors:** Esra Ay, Burak Eken, Tuğba Önal-Süzek

**Affiliations:** Department of Computer Engineering, Faculty of Engineering, Mugla Sitki Kocman University 48000 Mugla, Turkey

## Abstract

According to World Health Organization (WHO) 2016 report, there are over 650 million obese adults and more than 2 billion overweight individuals in the world and it is estimated that this number will reach 2.7 billion in 2025 [1]. A sedentary lifestyle with low physical activity is considered to be one of the most effective environmental effects leading to various chronic disease phenotypes such as obesity and metabolic syndrome. On average, every 1 out of 3 people over the age of 20 in Turkey are known to have struggled with the metabolic syndrome [2]. Our project aims to apply the concept of “serious gaming”, to entertain people, play games, socialize and exercise in parallel to increase the ratio of the healthy individuals in our society. In this project, we applied machine learning techniques to integrate real-life accelerometer and gyroscope sensor data obtained from mobile phones to develop an interactive mobile based exercise game which does not require any external device such as smart watches. To our knowledge and research, our game is the first mobile-only interactive serious game that integrates machine learning techniques and an encouraging virtual environment to the individuals in need of exercise.

## I. Introduction

Inadequate knowledge of the society about the long term consequences of physical activity for health implications and the adoption of an increasingly sedentary modern lifestyle had been one of the most important factors leading to higher incidences of chronic diseases such as obesity, cardiovascular diseases, hypertension, and diabetes. The negative effects of technology as well as the urbanization and scarcity of the suitable places for their physical activities have triggered physical inactivity. In this project we delved into this problem and to encourage people to exercise by developing a serious game to let users enjoy exercise without searching for a comfortable physical venue or companion. Serious games in healthcare are focused on four main areas; 1) games for rehabilitation providing a fun environment to improve patients’ cognitive and motor skills by using simulations and virtual reality environments during rehabilitation 2) games for health promotion and education raising awareness of the target population in aspects of diet, exercise, hygiene, and social abilities. 3) games for training physicians and healthcare professionals providing a simulated environment to reduce medical errors and subsequent healthcare costs[3,5] 4) games for distracting patients during painful procedures for focusing a patient’s attention away from the pain caused by their treatment with the famous example of the virtual reality game Street Luge shown as an effective and suitable pain relief mechanism[3,6]. The games for interactive exercise most similar to our product we found so far are “Pokemon Go”, “Zombies, Run!”, “Superhero Exercise”, “Walking”, “Geocaching”, “SpecTrek”, “BallStrike”, “Jump, Jump, Froggy” [4]. However the games listed in the four categories above and the other serious game examples we found either required an external device such as smart watch, a pen, a game-specialized virtual reality (VR) console or a connected computer all of which limit our main goal to reach-out to wide masses without any financial obligations like membership fees or equipment costs. Although the first FDA approved mobile game EndeavorRx [7] similarly uses the mobile device sensors, EndeavorRx specifically aims the cognitive improvement of the Attention Deficit Hyperactivity Disorder (ADHD) patients and does not aim to encourage an increase in the physical activity of the general population as our program.

## II. Methods

### A. Design Phase

Canvases and game screens prepared by each game are prepared by the authors. Game login and logout screens are created using the Unity launcher screen. The purpose of creating basic screens is to collect sensor data as well as check and test various ML algorithms in the game. The prototype screens required for the game were created by purchasing the required assets, a player avatar and a city simulation in which to exercise was created.

#### 1) Game Canvas

We designed our character and game map on the game canvas. The map we created was designed as a straight road going to infinity around a relaxing landscape (Fig. 1). Our character accompanies the user and runs on this road along with the mobile user for a certain time. For the map we prepared, assets were purchased from the unity asset store. Our character is taken from the unity asset store and placed in our game (Fig. 2).

**Fig. 1.**
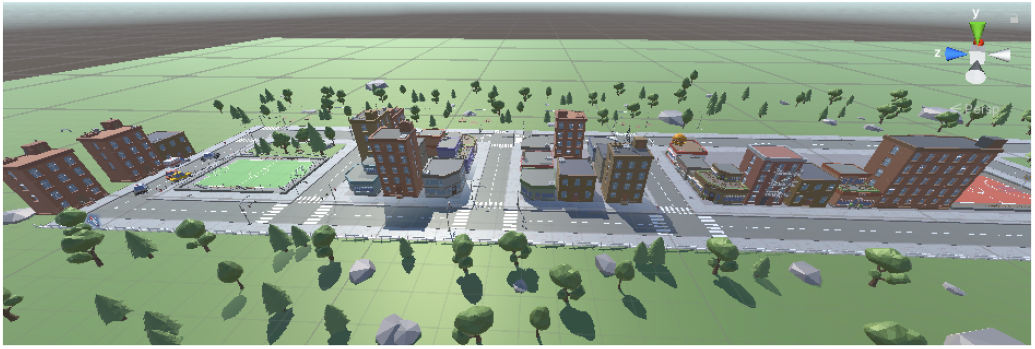
Map of game

**Fig. 2.**
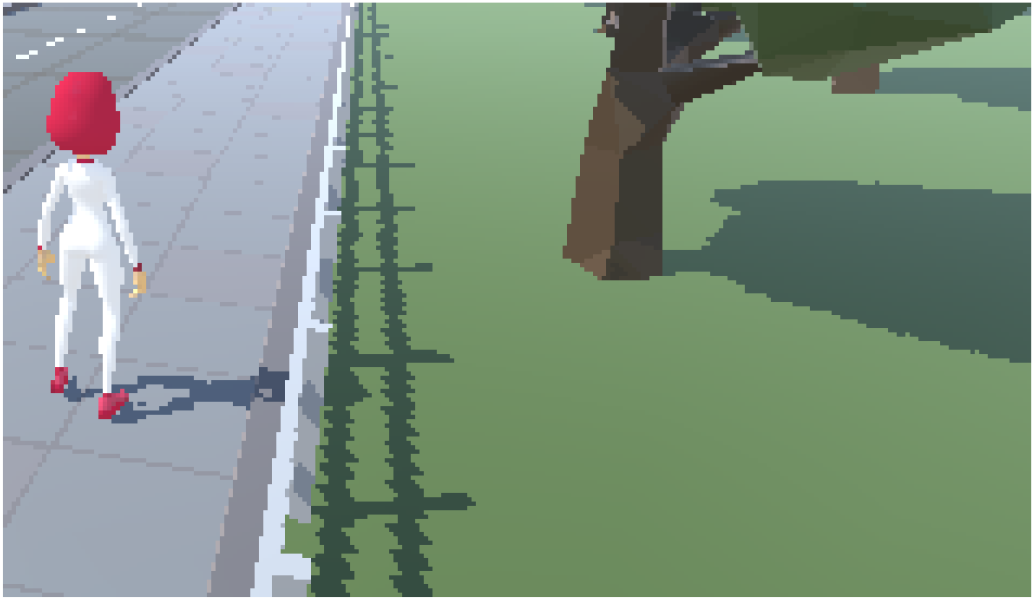
Player of game

Scale and rotation are adjusted while adding the player to the designed map. Light and camera settings have been defined for the player who will run backward. Thanks to these settings, the camera will follow the player and the player will be able to be displayed continuously on the game screen.

The animation controller panel has been created for animations that vary according to the player’s speed level (Fig. 3). In this way, the player whose speed level drops, can stop suddenly while running or start running while walking.

**Fig. 3.**
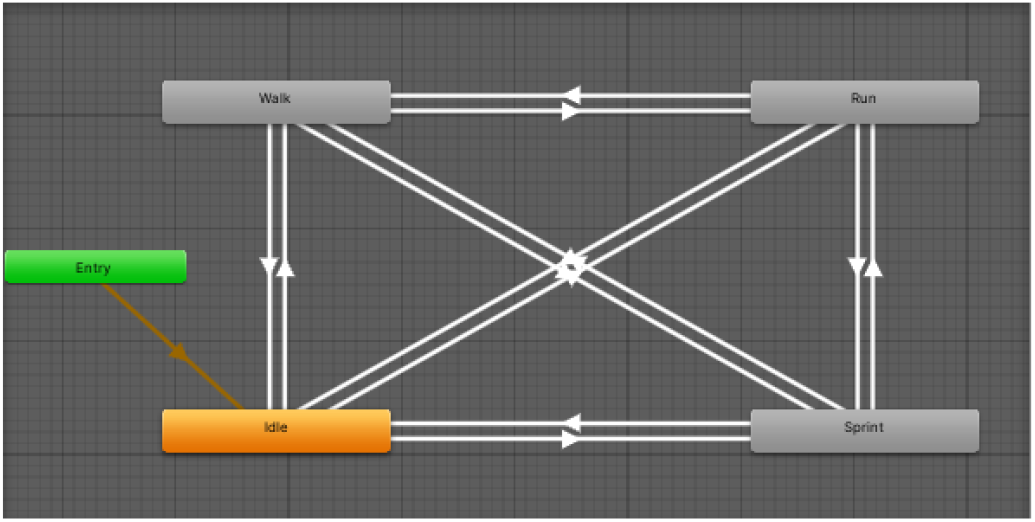
Controller Design

Animation controls for our character have been prepared and integrated into the player. Thus, our character will change its animation according to the speed of the user (Fig. 4).

**Fig. 4.**
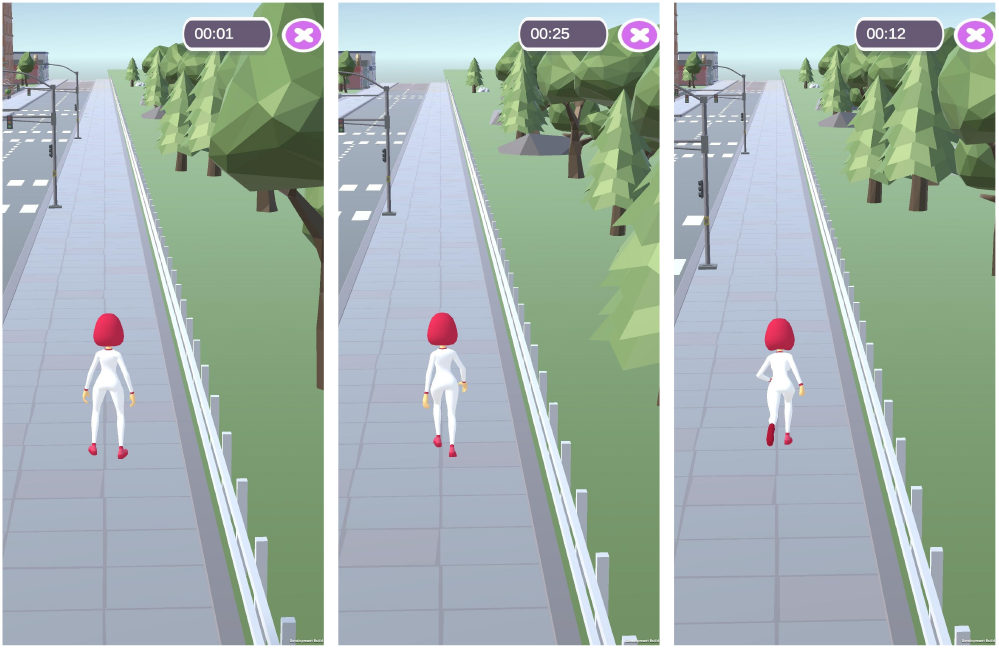
Player animations; IDLE, WALK and RUN

#### 2) Main Menu and Control The Sensors Canvas

Accelerometer and gyroscope are needed to collect data in the background of the game and predict the speed level. While the accelerometer is found in almost every device, the gyroscope is a rarer sensor. For this reason, whether the device has a gyroscope or not is checked on the home screen of the game. If the device does not have a gyroscope, the user is informed and the game is closed (Fig. 5).

**Fig. 5.**
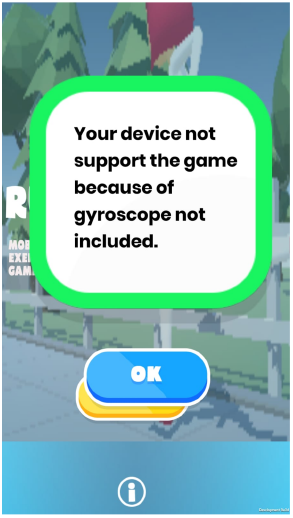
There is no gyroscope error

The name of the game and the inside of the game were used to design an image entry screen (Fig. 6). Added to meet the user in the game login screen in case of a gyroscope. Also, a button to launch the game and an information button that guides the user to an informative note indicating that our game is supported by TUBITAK 2209A program has been placed.

**Fig. 6.**
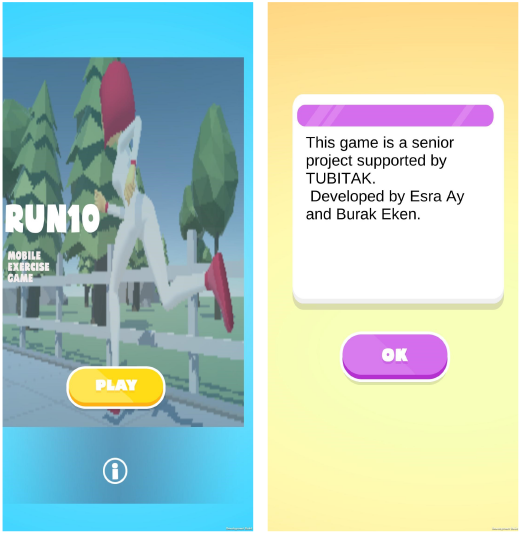
Main menu and Information about project

#### 3) Pause Game Canvas

During the game, we switch to this screen with the pause button in the upper right corner. Thanks to this screen, the user can return to the main menu, leave the game, resume the game, or restart (Fig. 7).

**Fig. 7.**
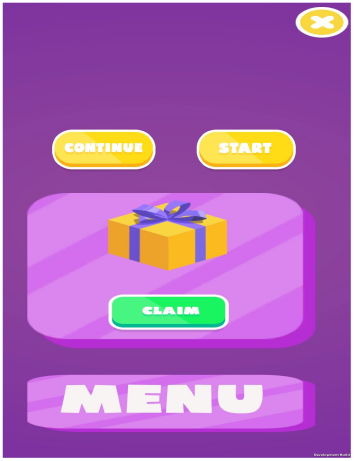
Pause screen

#### 4) End Game Canvas

At the end of the game, the user encounters this canvas (Fig. 8). On this screen, the user displays the score collected. There are also two buttons to return to the main menu and start the game again.

**Fig. 8.**
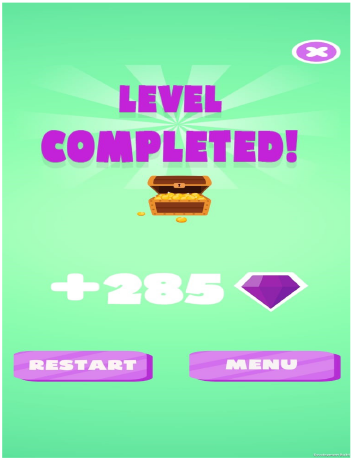
Game over screen

### B. Machine Learning Phase

Accelerometer and gyroscope are two main mobile sensors used in our game. Accelerometers in mobile phones are used to detect the orientation of the phone. The gyroscope, or gyro for short, adds an additional dimension to the information supplied by the accelerometer by tracking rotation or twist movements. An accelerometer measures the linear acceleration of movement, while a gyro on the other hand measures the angular rotational velocity. Both sensors measure the rate of change; they just measure the rate of change for different things. In practice, that means that an accelerometer will measure the directional movement of a device but will not be able to resolve its lateral orientation or tilt during that movement accurately unless a gyro is there to fill in that info.

Two different studies were carried out to determine the classification algorithm to be used in the game and to integrate it into the game. The first stage was implemented experimentally using Rstudio. Mobile sensor data were normalized and, trained using different classification algorithms, and their success rates were valuated.

After determining the algorithm with the best performance, a script was prepared to collect the sensor data within Unity. For Unity, the data set was created again as the sensor values received in Unity needed to be raw and Unity already normalized the input values. At this second stage of data collection, the mobile phone was connected via cable to the computer and data was collected for each speed level. The collected data were classified by passing through the KNN algorithm prepared in C# programming language using Collections, Generic, Linq, IO, Threading.Tasks libraries, and user testing was performed by printing the speed level on the screen.

#### 1) Experiments Using Mobile Sensor Recorder Application

In the experimental stage, gyroscope and accelerometer data to be used for classification were collected with an android sensor recorder application. While doing this, a treadmill and speed stages were used on it. Separate data were collected for each speed range on the 8-speed treadmill, and it was collected by moving the phone in the playing position at all speed stages. Then, a large data set was created by combining the data labeled according to each speed stage. RStudio was used for data classification and preparation. By combining all the obtained data into data frames, a function was written to normalize the data to calculate the average values of the physical activity performed in small time intervals as normalized data was required for the classification algorithms.

Classifications were implemented in RStudio using the created train dataset. Firstly, imputation was performed on the data and the dataset was split as train (70%) and test (30%). Three models based on KNN, Naive Bayes, and Decision Tree were prepared with the training dataset.

According to the accuracy and other results, successful classifications were made with the data set as can be seen in the graph Fig.9. Then prediction was made according to the models created using an independent test dataset. Although Naive Bayes is a successful model, the sensitivity value was found less than the rest of the algorithms. It is determined that the classifications that will best match our game are KNN and Decision Tree. All the classification codes are dumped as a report with RStudio in .rmd file format.

**Fig. 9.**
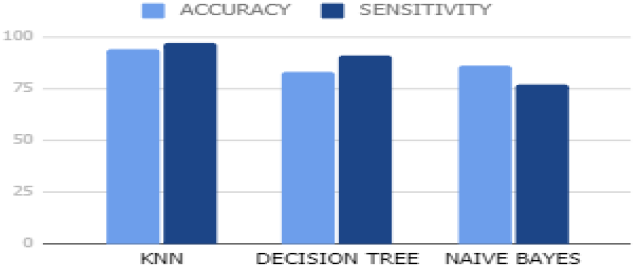
Accuracy and Sensitivity of Classification Algorithms

During classification, the normalized values, maximum-minimum values, and standard deviations of the collected data are taken into consideration. Features that are important in classification are created by processing the data collected with the Sensor Recorder. Classification is performed by analyzing the gravity and the accelerometer data. Some functions are very helpful in classification with their differences according to speed limits.

In the Variable Importance chart Fig.10, we observed that some features are more important than the others. This is due to the existence of specific value ranges and properties for each speed level.

**Fig. 10.**
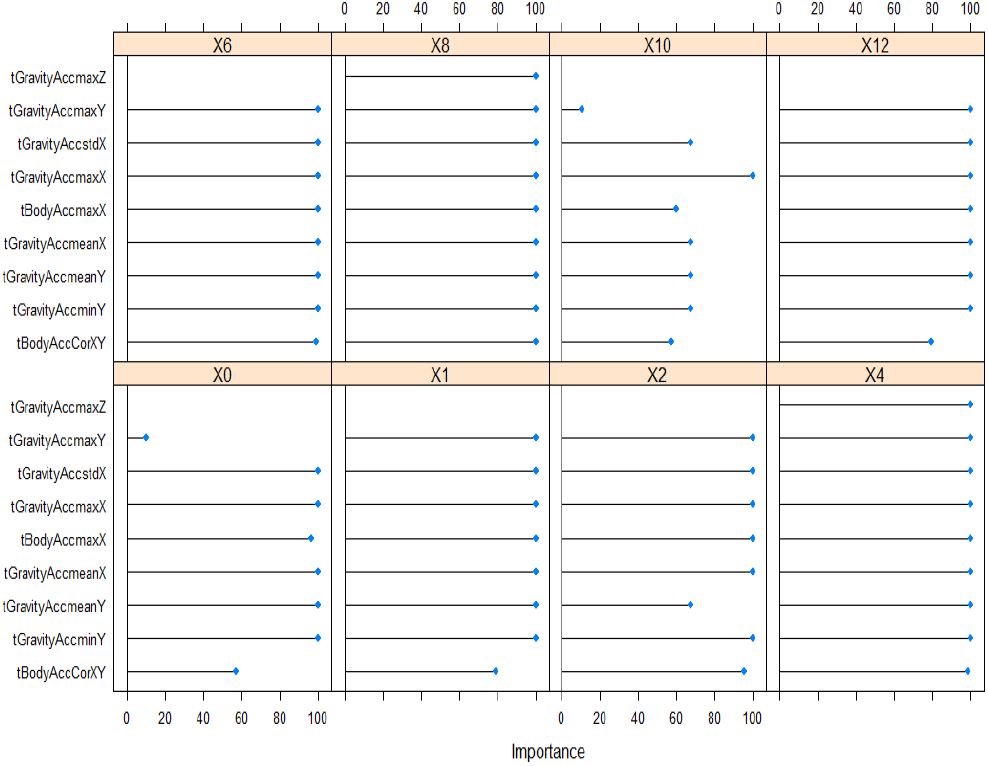
Variable Importance

When we examined further (Fig. 11), while the maximum value of acceleration in the Z-axis is an ineffective variable for many speed levels, it can be decisive for 8 and 12-speed levels. Acceleration, which is important for all velocity levels, takes place on the X-axis. When the maximum values and standard deviations of the values on the X-axis due to gravity and taken from the accelerometer are examined, we can say that they change in direct proportion to the velocity level.

**Fig. 11.**
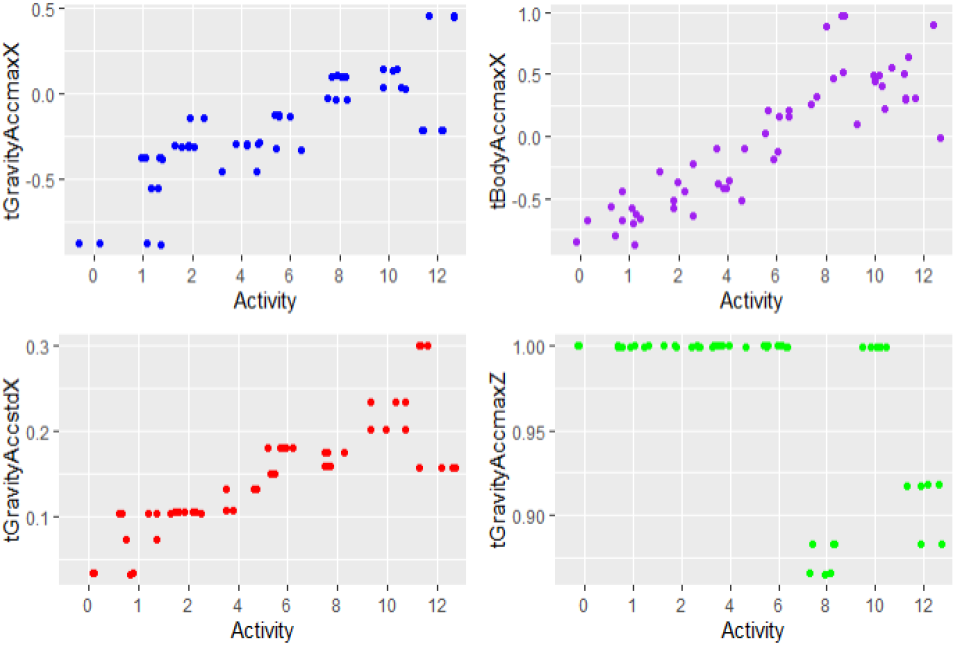
Analyzing Most Important Features

#### 2) Classification in Unity

This task was completed in Unity to predict the user’s movement speed. Same application has been re-implemented in Unity due to the data set difference in Unity. While this small application is running, it also creates the data set. The same treadmill and speed levels were used on the treadmill as in the original experiment. To collect the data the application is started on the computer connected to the mobile device with a cable, and the data received from the mobile device is tagged and recorded. To increase the actual performance of the game, the readymade plugins were omitted and several learning algorithms were compared. As a result of our research, the most suitable classification codes were determined [8]. All permissions and rights of the classification codes were obtained from Mr. Zevedei Lonut who prepared it. A c# script has been prepared for dynamically collecting and processing the sensor data that can classify the dynamically collected data according to the pre-prepared train data in the KNN algorithm. The KNN scenario has been integrated into the game.

## III. Results

As a result of the project, combining the machine learning and game programming techniques a unique serious game prototype is deceloped which can be played by anyone with access to a Android mobile device with accelerometer and gyroscope. With the budget provided by Tubitak 2209A program, missing device deficiencies are completed and several in-game assets are purchased yielding a better end-user experience. With the help of the classification algorithm, the user speed data set was reduced to 8 levels. Then we found out that determined speed level matched well with the in-game animation. As a result, a mobile game prototype has been successfully developed that synchronizes the real-time sensor data along with the incorporation of the remaining screens and proper detection of the right move of the player. As our main goal, we aimed to encourage the user for a more fun exercise environment where the user can self-improve gradually and achieved their own personal goals, and incorporate physical exercise as a part of daily life.

## IV. Conclusion

Our serious gaming project has the potential to minimize the lack of physical activity that has become a major problem in the western world. While doing this, our primary goal is to enable our users to exercise in the environment they prefer and thus encourage them for more exercise. Our game is available for all Android mobile devices accessible to any age audience. For future work, we are going to provide the user the option to select among multiple background environments and exercise in a multiplayer environment where the users will be able to exercise with their friends/partners living far away. Due to its easy accessibility through everyday mobile devices without any external hardware requirement, our mobile application has the potential to become a part of users’ daily life and increase the physical activity of the general population. As the next step of our prototype, our game will be extended to a multiplayer and multi language platform. After the multiplayer option is going to be deployed, the actual goal of our project of motivating the user’s physical activity by social interaction will be actualized, thus can be assessed objectively. For this reason, immediately after our multi-user option deployment, we are planning to formally assess our game’s effectiveness via user satisfaction survey.

## V. Software Availability

Our game APK is available at https://github.com/esraay/RunOn/tree/master/APK. It will be publicly available from Google Play upon acceptance of our manuscript.

## VI. Acknowledgement

We acknowledge the support of TUBITAK 2209-A University Students Research Projects Support Program grant that provided the funding to purchase the assets and the android device with the necessary sensors for our project.

